# Large cognitive fluctuations surrounding sleep in daily living

**DOI:** 10.1101/2020.06.18.159517

**Authors:** Reto Huber, Arko Ghosh

**Affiliations:** Child Development Center, University Children’s Hospital Zurich, Switzerland & Department of Child and Adolescent Psychiatry and Psychotherapy, Psychiatric Hospital University of Zurich, Switzerland; Institute of Psychology, Cognitive Psychology Unit, Leiden University, The Netherlands

## Abstract

It is well recognized that cognitive output fluctuates surrounding sleep. Such fluctuations are rarely investigated in subjects’ daily living, partially due to the inaccessibility of sleep laboratory technology and cognitive testing. Here we leverage the continuous and long-term smartphone touchscreen interaction logs to study the patterns of cognitive output surrounding sleep and contrasted these to the patterns underlying physical activity captured from wrist-worn actigraphy. According to spectral density analysis, both cognitive and physical activity was dominated by diurnal (∼24 h) and infra-radian (∼7 days) rhythms. However, these rhythms differed in a domain-specific manner. The proxy measures of cognitive performance – tapping speed, unlocking speed, and app locating speed – contained lower-powered diurnal rhythm than physical activity. Still, the amount of smartphone usage showed the strongest diurnal rhythm – even when compared to the ambient luminescence levels experienced according to actigraphy. Interestingly, the cognitive rhythms were not in sync with physical activity, as cognitive measures peaked later in the day and on weekdays rather than weekends. The difference between cognitive and physical activity became vivid during bedtime and subjects routinely interacted with their smartphones during the actigraphy labelled sleep period. The cognitive measures in this period were worse in comparison to the hour before or after sleep. Therefore, smartphones can seamlessly capture the dynamic fluctuations in cognitive output including during spontaneous awakenings. We conclude that the rhythms underlying cognitive activity in the real world are distinct from physical activity and this discord may be a hallmark of modern human behaviour.

## Introduction

We all experience that performance in daily tasks fluctuates strongly according to rhythms of different length. The best-studied rhythms in a laboratory setting are diurnal ^1,2^. Essentially, cognitive functioning across the 24 h depends on both circadian rhythms and sleep-wake duration-dependent fluctuations of sleep pressure. Specifically, the endogen circadian rhythm significantly impacts various performance measures in a ∼ 24 h rhythm, with performance being high during the day and deteriorating during the night-time ^3,4^. On the other hand, performance suffers also under high sleep pressure condition as a function of the duration of prior wakefulness ^3,5^. Additionally, sleep inertia, i.e. morning grogginess, reduces performance in the first 30 to 90 minutes after wake up ^6^. Ultimately the actual output of the system depends on the interplay between these different factors (circadian, sleep pressure, sleep inertia). We also know that misalignment of these internal processes which typically come along with extreme conditions (like shift work, jet lag or sleep restriction) can have dramatic consequences e.g. in terms of accidents and work-related errors ^7^. Though we have a substantial understanding of the fluctuations of cognitive functions based on the described factors, it is difficult to capture and study the consequences of these fluctuations in real life. Moreover, rhythms with a period longer than 24 hours are understudied because of the difficulty in capturing them in a laboratory setting. One promising way to overcome these empirical challenges is to leverage the daily digital interactions which may yield proxy measures of cognitive functions ^8–10^. According to one recent report leveraging the keypresses while on the web search engine, the speed of the keypresses fluctuates according to the time-of-the day akin to what has been found based on reaction time like tasks in the laboratory ^11^.

The unobtrusive sensing of cognitive performance in daily living conditions may provide some much-needed insights into the nature of the fluctuations under regular sleeping conditions i.e., without instructed sleep deprivation. Our understanding of these regular fluctuations is largely driven by the baseline measures obtained before sleep deprivation and other protocols in controlled laboratory settings^12^. For instance, sensorimotor cognition captured by using a psychomotor vigilance performance (PVP) test remains stable through the waking hours before the sleep deprivation but memory performance captured using a different test fluctuates through the day in the same period ^4,13^. The lack of data on cognitive fluctuations also percolates to the periods in daily living where sleep is fractured i.e., during the brief periods of interruptions in sleep due to either endogenous or exogenous reasons. This gap perhaps persists simply due to the unpredictability of these events – in particular when the interruptions are endogenously triggered - and thus out of reach of conventional performance assessments. Regardless, if people are woken up from their sleep to perform tasks akin to emergency response by a medical worker, executive functions, in particular, appear vulnerable ^14^.

In addition to the unavailability of the measures from particular times of the day, the generally sparse nature makes it difficult to estimate the precise nature of the rhythms that govern cognition. Towards this, the dominating periods must be estimated without imposing a priori waveform with a 24 h period. For instance, continuously measured physical activity (actigraphy) can be processed using spectral density analysis to establish the inter-individual differences in the near 24 h period, and these variations may be markers of clinical conditions ^15–17^. For cognitive measures, the fundamental issue of which period and the corresponding inter-individual differences remains unclear.

The ubiquitous smartphone interaction logs termed as tappigraphy can be converted into proxy measures of cognitive performance ^9^ and may provide the dense data needed to capture the fluctuations surrounding sleep. For instance, the tapping speed can be used as a proxy for cognitive processing speed that is conventionally based on reaction time tasks conducted in the laboratory or laboratory-like settings ^8^. Moreover, the near-continuous nature of tappigraphy will help detect a range of rhythms as done for actigraphy. In a recent study, we found that users typically engage on the smartphone through all the waking hours, and also into sleep to an extent that the longest smartphone usage gaps at night provide a reliable proxy for actigraphy estimated sleep ^18^. Interestingly, virtually all the users engaged on the smartphone – even if briefly – during the actigraphy estimated sleep ^18^. This phenomenon of sleep fracture, in particular, opens the opportunity to measure the cognitive status during the period of sleep fragmentation by using tappigraphy.

In this study we defined three different tappigraphy parameters to ubiquitously capture the cognitive processes akin to that measured in reaction experiments in the laboratory: (a) the time that is taken to go from one touch to another, (b) the time that is taken to unlock the phone and (c) the time that is taken to locate app icons on the home screen before launch. This yielded a time series of measurements enabling spectral density analysis of the cognitive fluctuations to identify the oscillations that dominate the cognitive outputs. We contrasted these measures to actigraphy (including ambient luminescence) captured using a wrist-worn wearable. Finally, through the combination of actigraphy and tappigraphy, we estimated the cognitive status during the actigraphy labelled periods of pre-bed, ‘sleep’, and rise times.

## Results

### Periodicity in luminescence, physical activity, and tappigraphy

The wearable and tappigraphy signals fluctuated through the recording period (Fig. 1). To quantify the periodicity of these fluctuations, we used the Lomb-Scargle method. The population average traces were used to quantify the consistency of the behavioural patterns in the sampled population. The periodograms revealed a consistent ∼24 h periodicity across all of the signals and a less prominent ∼7 d periodicity for the ambient luminescence, physical activity, smartphone usage (no. of touches), and the two proxy measures of cognitive processing speed: tapping speed (TS) and unlocking speed (US) (Fig. 2).

**Figure 1.**
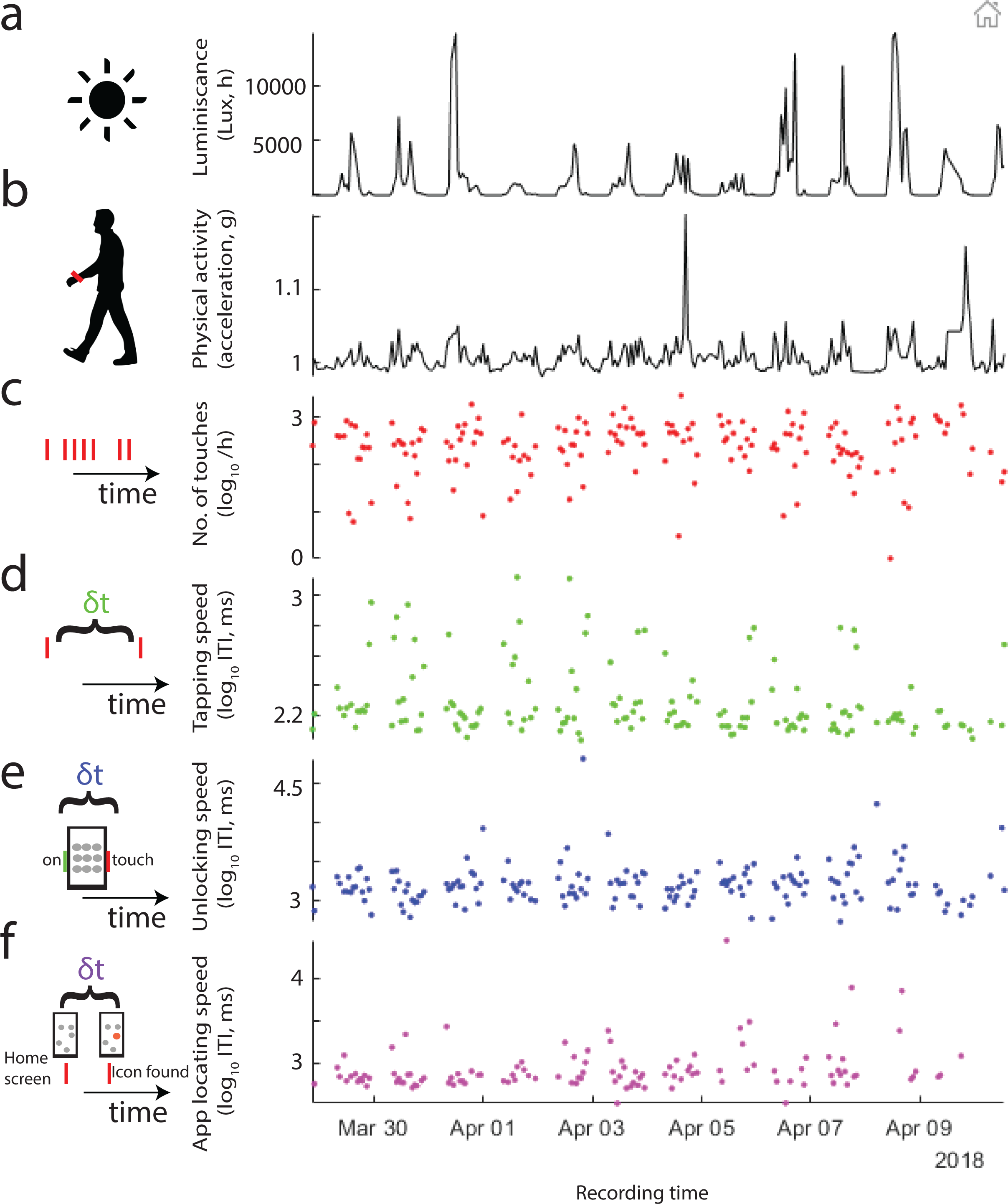
The range of measures captured in daily living conditions in this study. Wearable (a-b) and smartphone data from one exemplary subject monitored over 21 days. (a) The amount of ambient light was captured on the wrist along with (b) the acceleration used here as a proxy for physical activity. We quantified the smartphone behaviour in hour-long bins in terms of (c) the number of touches, (d) the speed of the interactions measured as the interval between subsequent touches (fastest 25^th^ percentile at each bin, tapping speed, TS), (e) the time taken to unlock the screen (unlocking speed, US) and (f) the time to select an app icon while on the home screen (app locating speed, ALS).

**Figure 2.**
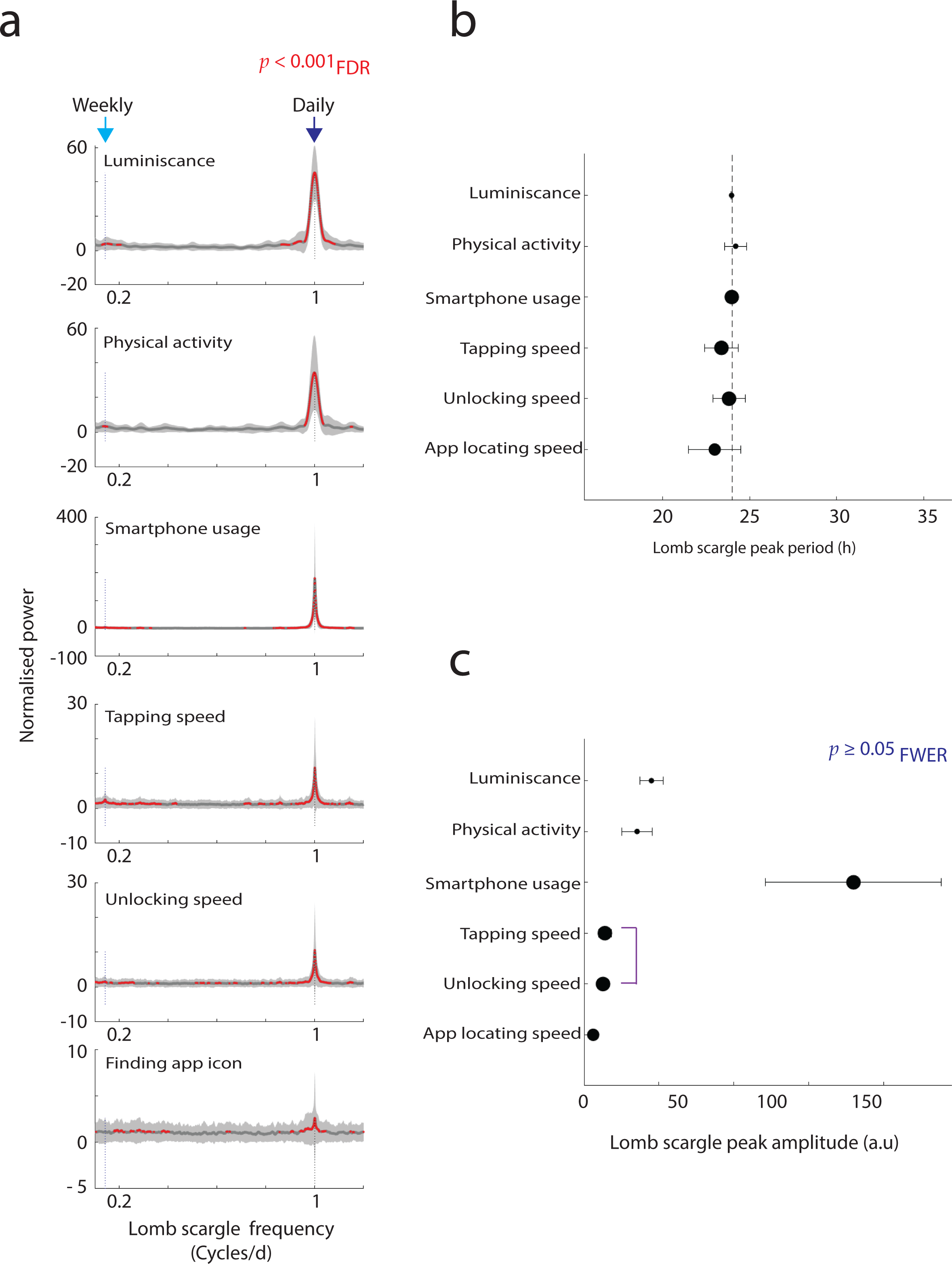
Lomb-Scargle periodogram reveals the periods that dominate the wearable (luminescence and physical activity) and smartphone parameters. (a) Mean periodogram and their corresponding confidence intervals (95%), with significant differences from zero amplitude signal marked in red. (b) Mean peak periods derived from the periodograms and their corresponding confidence intervals. (c) Mean peak normalised powers and their corresponding confidence intervals. The size of the shapes in b & c correspond to the sample size (see main text for the corresponding degrees of freedom).

By using the periodogram peaks we next estimated which period consistently dominates the signals. First, we contrasted the location of the periodogram peaks with 24 h periods using one sample t-tests to establish deviations from this anticipate period (α_FWER_ = 0.0083). The mean peak periodicity of ambient light fluctuations was 23.96 (*p* = 0.002, *t*(69) = -3.27), for movements it was 24.19 (*p* = 0.22, *t*(70) = 1.23), for smartphone usage it was 23.97 (*p* = 4.24 x 10 ^-6^, *t*(183) = -4.74), for TS 23.38 (*p* = 0.01, *t*(184) = -2.57), for US it was 23.82 (*p* = 0.43, *t*(184) = -0.78) and for ALS it was 23.00 (*p* = 0.01, *t*(156) = - 2.65). Next, we compared the peak locations across the signals to find that the periods were domain dependent (Fig. 2, *p* = 0.01, *f*(5,846) = 3.01, ANOVA).

There were pronounced inter-parameter differences between the power estimates (i.e., in the normalised peak amplitudes of the periodogram, *p* = 7.22 x 10^−74^, *f*(5,848) = 87.02, ANOVA, Fig. 2c). The power of smartphone usage showed the highest ∼ 24 h peaks relative to any of the signals and the ALS showed the weakest peaks. Notably, the proxy measures of cognitive processing (TS, US, and ALS) speed show a lower amplitude than the wearable measures of luminescence or physical activity (Fig. 2).

### Time-of-the-day effects in physical and cognitive signals

Upon establishing the dominance of ∼ 24 h rhythms in the physical and cognitive fluctuations we next focused on how the gathered measures relate to the time-of-the-day. As anticipated from the periodograms, the central tendencies revealed substantial time-of-the-day fluctuations across the parameters (Fig. 3). To systematically address at which hour the performance consistently peaked we relied upon cosinor analysis with a fixed 24 h waveform (with individual fits corresponding to *p* < 0.05). The hour at which performance peaked (the cosinor acrophase) depended on the parameter (*p* = 0, *f*(5,673) = 25.64, Watson-Williams multi-sample test). Follow-up tests revealed that the peak for ambient light exposure preceded all other measures, whereas the peak for unlocking speed lagged all other measures except for ALS (Fig. 3). Although the hour of peak performance for the proxy measures of cognitive processing occurred between 16 to 17 h, the differences between the peak vs. off-peak performance was the most pronounced for TS (*p* = 3.23 x 10^−29^, *t*(125) = 14.78, paired t-test), followed by US (*p* = 3.48 x 10^−12^, *t*(117) = 7.76) and the ALS (*p* = 0.0062, *t*(39) = 2.89).

**Figure 3.**
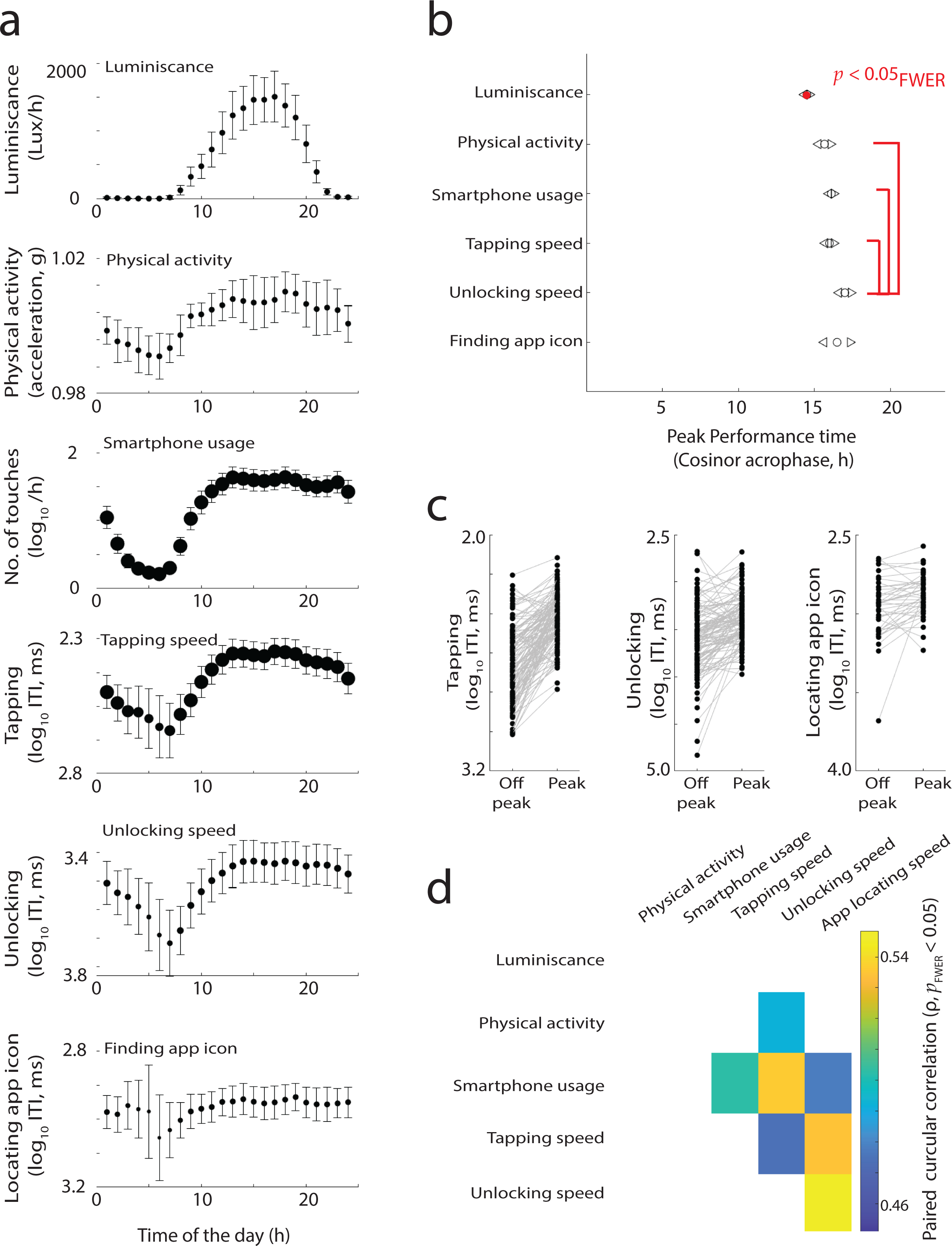
Time of the day reflects on physical activity and processing speeds captured on the smartphone. (a) Time of the day fluctuations in mean values and the corresponding 95% confidence intervals. (b) Cosinor fits revealed the peak performance times on the 24 h clock, the confidence intervals marked with triangles. (c) The comparison of processing at the acrophase vs. off phase in the sample, with each individual represented with a connecting line. (d) The inter-individual differences in the time of peak performance (cosinor, acrophase) are related to each other. The circular correlation coefficient is shown for the significant relationships.

Next, we addressed whether the inter-individual differences in the acrophase were correlated across parameters (Fig. 3). In particular, we were interested in the putative determinants of the cognitive processing speed. We used paired circular correlations to address these relationships. Interestingly, the subtle variations in the ambient light acrophase were not correlated to any of the measures. However, physical activity was correlated to only one of the cognitive measures: US. Furthermore, smartphone usage was related to all measures of the cognitive processing speed.

Similarly, we also followed-up on the ∼7-day rhythm identified using the periodogram based on time-of-the week analysis to find systematic variance according to the day-of-the-week, for all measures except the US and ALS (Supplementary Fig.). The day on which the signals peaked varied according to the measured parameter (*p* = 0, *f*(5,838) = 52.77, Watson-Williams multi-sample test). Furthermore, while physical activity and luminescence peaked around the weekend, smartphone usage and tapping speed peaked around the weekday.

### Poor cognitive performance during actigraphy labelled ‘sleep’

The results from the time-of-the-day effects revealed that there were smartphone interactions captured at all hours of the day in the sampled population. Struck by this phenomenon we set out to explore how the cognitive performance fluctuates surrounding sleep and, more uniquely, while in actigraphy defined sleep (i.e., while lying still in bed and using the smartphone and yet classified as sleep by the Cole–Kripke algorithm on actigraphy). From each individual, we pooled all the instances of usage at three different periods: *pre-bed* defined as 1 h immediately preceding actigraphy defined sleep time, *bed* defined as during the putative sleep (actigraphy defined) time and *rise* defined as 1 h immediately following the sleep period (Fig. 4). We focused on the subset of the population where the parameter estimation requirements were satisfied to yield a measure in each of these periods. TS fluctuated only marginally across the three periods (*p* = 0.07, *f*(2,65) = 2.73, ANOVA). Follow-up paired t-tests revealed marginal slowing in *bed* vs. *pre-bed* (p = 0.05, t(65) = -2.0), and the *bed* vs. *rise* (*p* = 0.05, *t*(65) = 2.04). Furthermore, there was no difference between the *pre-bed* vs. *rise* (*p* = 0.59, *t*(65) = -0.54). US fluctuated substantially through these periods (*p* = 5.62 x 10-9, *f*(2,62) = 22.23), and the follow-up t-tests revealed a similar pattern as for TS albeit more exaggerated. The *bed* period compared to *pre-bed* was substantially slower (*p* = 5.01 x 10^−7^, *t*(62) = -5.6), and the *bed* period was also slower vs. *rise* period (*p* = 6.12 x 10^−5^, *t*(62) = 4.3). There was a marginal difference between the *pre-bed* and *rise* period, with the *rise* being slower (*p* = 0.03, *t*(62) =-2.20). ALS too fluctuated through these periods (*p* = 5.50 x 10^−30^, *f*(2,47) = 150.08). Interestingly, the *bed* period was no different vs. the *pre-bed* period (*p* = 0.52, *t*(47) = -0.64). However, the *rise* period was faster than both *pre-bed* (*p* = 0, *t*(47) = 12.98) and *bed* periods (*p* = 0, *t*(47) = 15.53).

**Figure 4.**
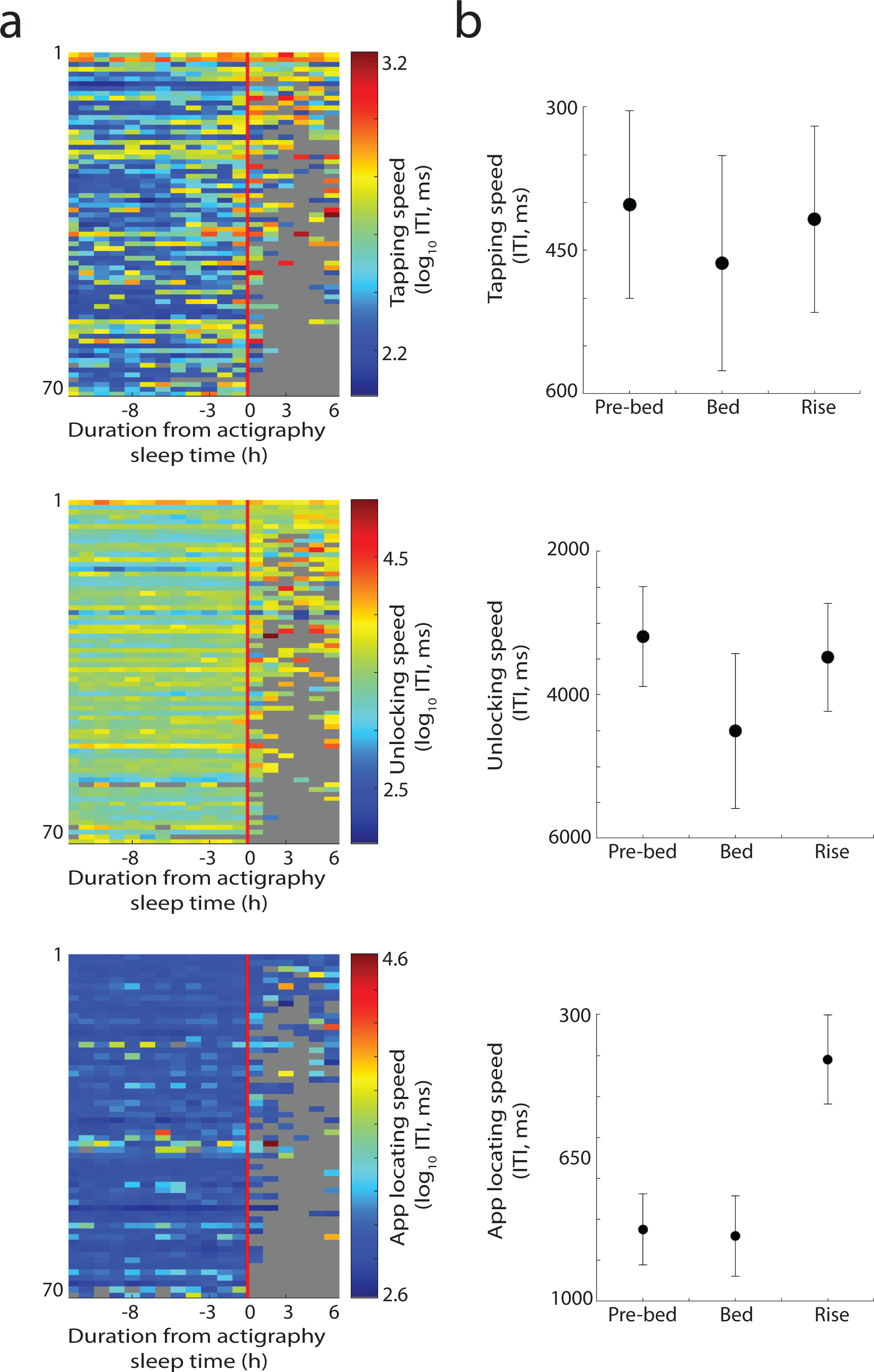
Cognitive processing speed captured on the smartphone during actigraphy labelled sleep. (a) The median processing speeds – in terms of tapping speed, unlocking time and locating the app icon - captured for each individual accumulated over the observation period. (b) The differences in mean processing speed captured during actigraphy estimated sleep in contrast to the values accumulated in the hour before sleep and after sleep (95% confidence interval). The size of the shapes in b correspond to the sample size (see the main text for the corresponding degrees of freedom).

## Discussion

The proxy measures of cognitive processes captured using tappigraphy revealed a range of systematic fluctuations and here we contrasted these to ambient luminescence and physical activity. The tappigraphy measures – like luminescence and physical activity – was dominated by ∼ 24 h rhythm. But there were visible differences between the cognitive and the non-cognitive rhythms in terms of the exact period, the power and even the time-of-the day when they peak. Some of these differences also extended to the ∼ 7 d rhythm. Intriguingly, the tappigraphy measures also allowed us to assess performance at odd hours including when in bed revealing a distinctly slow cognitive output.

As continuous measures related to cognitive output are mostly unexplored the general patterns of the signals observed in tappigraphy are of interest. The periodograms of the three proxy measures of cognitive processing speed (TS, US, ALS) revealed dominant powers at frequencies with periods of ∼ 24 h and ∼ 7 d. While all of the measures were dominated by 24 h cycles, there were substantial inter-individual differences and the exact period varied according to the parameter. This is in line with previous observations demonstrating deviations from the 24 h period in cognitively rooted performance parameters such as handgrip strength ^19^. We also observed ∼ 7 d in the cognitive parameters of smartphone usage and TS. This supports the idea that the time information – both in terms of time-of-the-day and time-of-the-week – is encoded in cognitive output ^20^.

Luminescence is a primary *zeitgeber* for circadian physiological rhythms according to observations mostly in the sleep laboratory ^21–23^. Still, cognitive processes may not be faithfully tethered to this in the real world. First, the cognitive processing measures were less dominated by ∼24 h rhythms in comparison to the luminescence captured at the wrist. Secondly, according to the fixed (24 h), cosinor analysis all of the proxy measures of cognitive processing peaked (became faster) later in the day in comparison to the experienced luminescence. Finally, population-level variance in tappigraphy time to peak was unrelated to the variance in the experienced luminescence. We speculate that cognitive processes follow diurnal rhythms that are partially independent of the experienced luminescence.

The processes that drive the well-documented rhythmicity in overall physical activity may not entirely overlap with the oscillators underlying cognitive activity in the real world. Towards this, cosinor analysis revealed some important separations between cognitive and physical activity. The US peaked later than physical activity and the variations in time to peak in physical activity were only related to one of the tappigraphy measures – US. The differences between physical and cognitive activity were further widened in the ∼ 7 d cycles and while tappigraphy measures (smartphone usage & TS) peaked during weekdays, physical activity (and experienced luminescence) peaked during weekends.

Interestingly, smartphone behaviour showed the strongest ∼ 24 h power (signal normalised) in comparison to the other measures considered here. This underscores the habitual nature of smartphone behaviour where it may be more driven by daily cycles than overall physical activity or the amount of light exposure. Nevertheless, the daily cycles were less powerful for the proxy measures of cognitive output suggesting discord between engaging in smartphone behaviour and the cognitive processing speed proxied on the smartphone. This discord was vivid in the time-of-the-day analysis and by using the fixed 24-hour cosinor analysis we found that UT peaked (was the fastest) later in the day in comparison to smartphone usage. This raises the possibility that the need or desire to engage on the smartphone and certain cognitive performance abilities are out of sync – i.e., phase shifted.

By leveraging smartphone touchscreen behaviour, we could sample cognitive fluctuations at the grey zone between sleep and wakefulness. People spontaneously interacted with their smartphones in the actigraphy labelled sleep periods and we leveraged these interactions to address the cognitive status in this ‘sleep’ period in comparison to the performance 1 h before and after this period. Now, admittedly actigraphy can overestimate sleep and people may engage on their smartphones while at rest in bed ^18^. Still, this provided us with an opportunity to assess cognition in this period of sleep fracture. Across the different proxy measures of cognitive processing, the performance was poor in this obscure period. The mechanism underlying this as in sleep inertia vs. pressure could not be clarified without polysomnography; it is possible the participants intermittently woke-up from sleep in the bed (inducing inertia) and it is equally possible they may have remained still without sleep (building sleep pressure). The current pattern suggests a dual contribution. In the hour after sleep, inertia can be considered to be maximal and yet the performance at sleep fracture was worse than this period. This suggests that an additional factor – such as sleep pressure – is compounded with sleep inertia to additionally degrade cognitive output in the obscure sleep period. Conversely, in the hour before sleep, sleep pressure can be considered to be maximal and yet TS and US degraded further at sleep fracture. Interestingly, ALS did not degrade further and perhaps the underlying processes are particularly sensitive to pressure rather than inertia. Such specific variations or lack there off are in keeping with the general notion that sleep impacts cognitive processes in a domain-specific manner ^24,25^.

Our approach of assessing cognitive fluctuations surrounding sleep in daily living conditions raises some important questions. First, we observed different rhythms in cognitive vs. the physical activity measures including the presumable *zeitgeber* of ambient light. It was not clear if these asynchronies were introduced by smartphone behaviour or if they are intrinsic properties captured on the smartphone. On a related note, the consequences of the differences between smartphone usage vs. the proxy measures of cognitive processing too need further exploration. These asynchronies may have important consequences for mental and physical well-being ^26^. Second, there is much to be addressed on why and how people behave at physical rest while in bed (i.e., actigraphy defined sleep). What are the cognitive and behavioural processes underlying these behaviours that spontaneously occur so close to sleep? Finally, unlike traditional cognitive testing, the parameters extracted from spontaneous smartphone behaviour do not allow us to specify the cognitive processes with precession even if they are correlated to conventional reaction time or even if they intuitively correspond to cognitive processing speed. Nevertheless, meaningful cognitive processes are inherently complicated and the approach of tappigraphy and the findings presented here is a key step to help unravel that complexity.

## Methods

### Participants

A total of 189 individuals were sufficiently sampled (with 235 addressing the recruitment call, full data is made available and see below). To be included in the study the subjects self-reported that they were healthy and without any ongoing neurological disease or medication. The data collection and analysis were approved by the ethical committees of Leiden University (Psychology Research Ethics Committee) and the medical ethics committee of Arnhem-Nijmegen. All subjects provided informed consent for the study. The age (reported by 165 participants) was a median of 23 years (min, 16 and max, 45).

### Actigraphy measurement

Actigraphy measures were obtained from a subset of participants (n = 79) and reported in a previous study ^18^. Participants wore GENEACTIV watches (Activinsights, Cambridgeshire, UK) on both the wrists, but only the measures from the left wrist were used here. The watches measured the 3-axis accelerometry along with the ambient luminesce and near body temperature, but only the former two measures were used here. The participants were instructed to wear the watches for 3 weeks continuously and this yielded measure lasting for a median of 21 days (min, 7 and max, 32). The 3 axis accelerometry was reduced by using *M* = √(*x*^2^ + *y*^2^ + *z*^2^), where M is the value used here and x, y and z correspond to the accelerations on the distinct axis. The Cole–Kripke algorithm was used to label sleep periods based on these measures as described in detail elsewhere along with the corresponding MATLAB codes ^18,27^.

### Smartphone measurements

The timestamp of touchscreen interactions and the corresponding app labels (as in Facebook, Launcher screen, Weather) were recorded using an app running in the background of the user’s device (TapCounter, QuantActions, Lausanne, Switzerland). Based on this labelled time-series of events the following parameters were estimated in hourly bins: (a) Smartphone usage, in the form of a number of touchscreen interactions in each bin while the phone was in an unlocked state, (b) tapping speed, in the form of the 25th percentile inter-touch interval accumulated from all of the screen ON sessions in each bin, (c) unlocking speed, in the form of 25th percentile inter-event interval between the two intervals, one, the touchscreen turning ON and two, the touch on the unlocked screen and, (d) app locating speed, as in the inter-touch interval between two consecutive touches on the home/launch screen (identified using the corresponding app label) before the launching of any app. As with the previous measures the 25^th^ percentile of the intervals in each hour bin was recorded. All of the smartphone parameters were transformed by using *log*_10_.

### Estimating the periodogram and the corresponding metrics

Lomb-Scargle periodograms were estimated (MATLAB, Mathworks, Natick, USA) and the power was scaled by the input variance. The periodogram was estimated between 0.05 and 12 cycles per day with a step of 0.001 cycles. The statistical significance (α = 0.001) of the power fluctuations were estimated against 0 using t-tests (LIMO EEG^28^) and multiple comparisons corrected using the false discovery rate (FDR, also on LIMO EEG). Inputs spanning longer than 10 days were used for this analysis. To compare the periodogram peaks at ∼1 cycle per day across the different smartphone and wearable parameters, the peak was determined within the range of 0.7 and 1.6 cycles per day. First, the peaks from the different measures were compared using one-way ANOVA (MATLAB, MathWorks, Natick, USA). These were followed-up with t-tests comparing all possible pairs of measures. The tests were corrected using Bonferroni correction of Family-Wise Error Rate (FWER, α = 0.05, Victor Martinez’s Multiple Testing Toolbox as implemented MATLAB)^29^. The 95% confidence intervals were estimated using the inverse of Student’s T cumulative distribution function (MATLAB). Follow up t-tests after ANOVA to compare the periodogram peaks (location and amplitude) across the different parameters were also corrected using FWER. Inputs spanning longer than 7 days were used for this analysis block focused on ∼1 cycle per day rhythm.

### Finding peak-of-performance in time-of-the-day using cosinor analysis

The acrophase of the sine wave fits obtained using Cosinor.m (implemented in MATLAB by Casey Cox)^30^. Inputs spanning longer than 7 days were used for this analysis. The time-of-the day fluctuations were compared across the different parameters using the Parametric Watson-Williams multi-sample test (Circular Statistics Toolbox for MATLAB) and as a follow-up, the same test was used in pairs. The inter-individual differences in the acrophase across the different parameters were tested for correlation using circular correlation (Circular Statistics Toolbox for MATLAB) ^31^. The statistical output was corrected for multiple comparisons using the Bonferroni correction of Family-Wise Error Rate (FWER, α = 0.05). The 95.0% confidence intervals were estimated using the same toolbox.

### Finding peak-of performance in day-of-the-week

The hourly smartphone and wearable parameters as described above were sorted according to the day of the week. Inputs spanning longer than 10 days were used for this analysis. The mean value from each day of the week was used to derive the location of the peak. These locations were converted into radians towards circular mean and confidence intervals (95%). The measures were compared for day-of-the-week differences across the different parameters as stated above for time-of-the day analysis, that is by using the Parametric Watson-Williams multi-sample test and follow-up paired tests were corrected for FWER.

### Estimating performance surrounding sleep

The hourly smartphone parameters were time-locked to the sleep times estimated using the Cole-Kripke algorithm on the actigraphy measures from the left wrist ^27^. The median values in the hour bin preceding, during and after the sleep period was estimated from each individual. To address if there were differences between these three measures were contrasted using two-way ANOVA (MATLAB, MathWorks, Natick, and α = 0.05).

### Code availability

All of the codes that were programmed for this study are made available at the dataverse.nl repository (<u>temporary repository link for peer-review</u>).

### Data availability

A reduced and de-identified data at 1 h bins are made available at the dataverse.nl repository. (<u>temporary repository link for peer-review</u>)

## Acknowledgements

We thank the students at the Applied Cognitive Psychology master’s program at Leiden University for aiding in the data collection. We thank Andrew Westbrook for sharing the smartphone data collected at The Donders Institute. We thank Leonardo Cohen for his advice during the preparation of this manuscript and suggesting the title.

## Conflict of interest

Arko Ghosh is a co-founder of QuantActions Ltd, Lausanne, Switzerland. This company focuses on converting smartphone taps to mental health indicators. Software and data collection services from QuantActions were used to monitor smartphone activity.

**Supplementary Figure.**
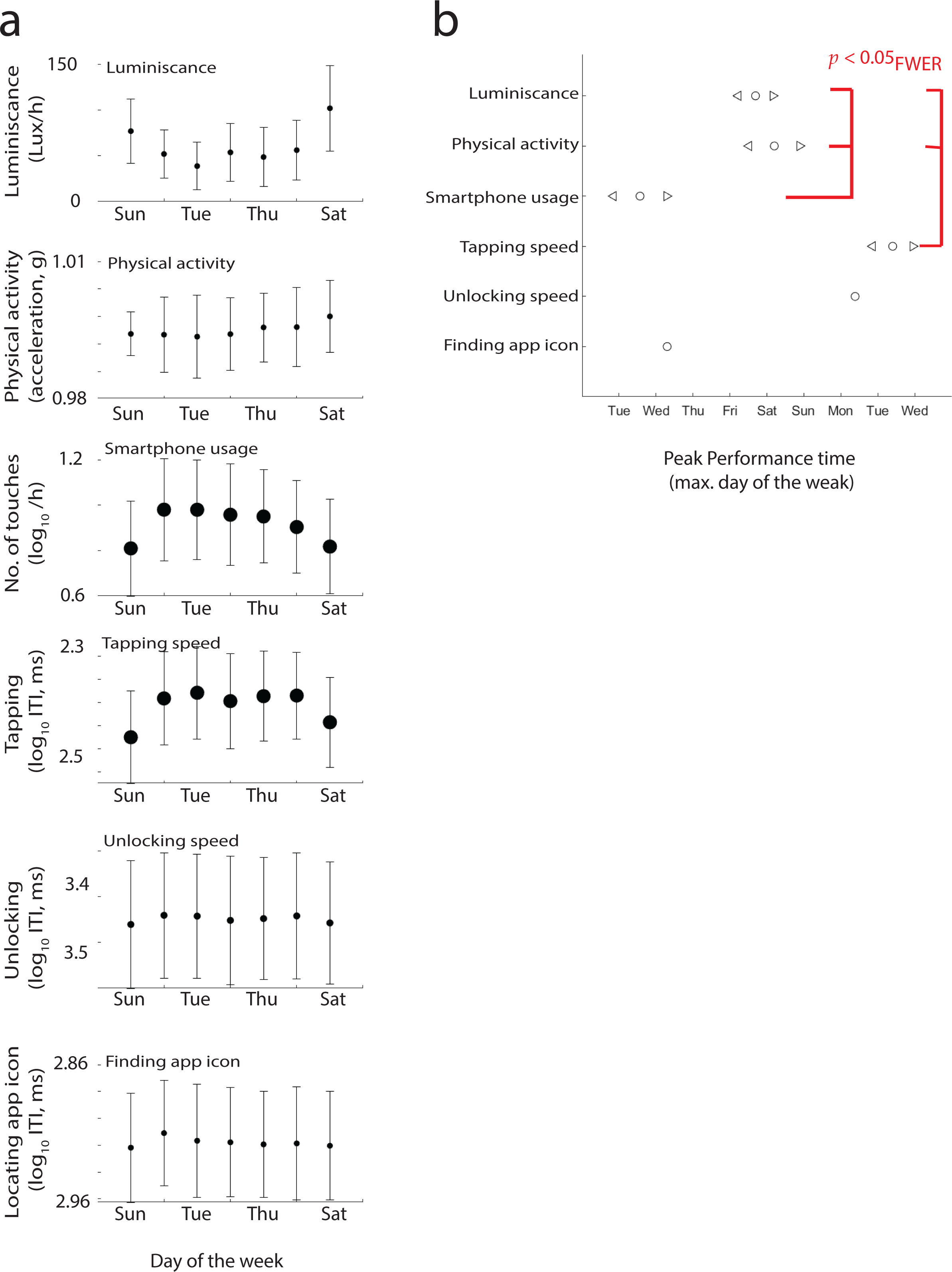
Day of the week reflects on physical activity and processing speeds captured on the smartphone. (a) Mean values and the corresponding confidence intervals (95%). (b) The peak performance in terms of the best (mean) performing day of the week and corresponding confidence intervals.

